# Cutting tree branches to pick OTUs: a novel method of provisional species delimitation

**DOI:** 10.1101/419887

**Authors:** Mikula Ondřej

## Abstract

Delimitation of species is crucial in all studies of biodiversity, its geographic patterns and evolutionary dynamics as well as in the corresponding conservation applications. In practice, operational taxonomic units (OTUs) are often used as provisional surrogates of the species, whose evidence-based and robust delimitation requires too extensive data and complex analyses. The novel method for this provisional species delimitation is suggested, which uses any phylogenetic tree with meaningful branch lengths as an input and delimits OTUs on it by identification of branches whose removal significantly changes structure of the tree. Such branches are considered to reflect interspecific differentiation that is assumed generally more erratic than intraspecific branching. It is called branch-cutting method as it evaluates structural importance of the branch by its cutting (shrinking to zero length) and inspecting impact of this operation on the average pairwise distances between tree tips. Tree tips can be also constrained to be either conspecific or heterospecific which allows the method to achieve more robust and informed delimitations and to focus on particular phylogenetic scale. Usefulness of the method is demonstrated on four empirical examples and comparison with similar methods is performed.

## Introduction

In many ecological and evolutionary studies species play a crucial role of fundamental units whose delimitation can affect all downstream analyses (Zachos 2016, Ch. 7). Yet doubtful species boundaries are commonplace and because time and resources necessary for their investigation are limited, it is upon the researcher to resolve them, at least provisionally. Such *ad hoc* delimited species may be also denoted as operational taxonomic units (OTUs). Following the massive spread of molecular techniques, the genetic data analysis became a cornerstone of virtually all ‘species delimitation’ and ‘OTU picking’ methods (Sites & Marshall 2003; Carstens et al. 2013). Some of these methods use explicit population genetic or evolutionary models, most importantly Hardy-Weinberg equilibrium of allele frequencies (Guillot et al. 2005; Huelsenbeck et al. 2011) and multispecies coalescent process explaining distribution of gene genealogies and ultimately sequence variation (Yang & Rannala 2010, Jones et al. 2015). Others treat species delimitation as a pattern-recognition problem, assuming that speciation leaves a specific signature in molecular data, distinguishable from intraspecific variation. Typically, genetic distances within species are assumed to be markedly lower then among species and distribution of genetic distances is therefore multimodal. It implies there is a characteristic difference between intra- and interspecific variation (so called ‘barcoding gap’), which may be estimated to establish OTU boundaries (Puillandre et al. 2012).

## The branch-cutting method

### Algorithm

Here, I present a novel OTU-picking method from the ‘pattern-recognition’ family. It operates on a single rooted or unrooted gene tree and considers *structurally important branches* as signatures of speciation. The basic assumption is that speciation is slower and less regular than mutation and thus it manifests itself by branches whose removal significantly changes structure of the tree. In other words it assumes each speciation to be so unique that its associated branches are not interchangeable in the overall picture of evolutionary process represented by the gene tree. When one sequence is replaced by another randomly picked from the same species, the structure of the tree is expected to be virtually unchanged and along the same line of argument shrinking of one intra-specific branch to zero changes little in the structure of the tree as the species is still well represented by its remaining representatives. This is not expected to be the case when inter-specific branch is removed. The structural importance (non-interchangeability) of a branch can be quantified as a decrease of the mean pairwise distance between tree tips after removal of the branch or, equivalently, after setting its length to zero. The branches are then ranked in increasing order of this loss score and just a few of them with the top ranks are retained. The length of all other branches, that are not retained, is set to zero which causes groups of tips to collapse into clusters where they are separated from each other by zero distance. These clusters are then considered as OTUs, i.e. provisionally delimited species. Metaphorically, the tree branches are cut in order to pick OTUs and thus the procedure is called *branch-cutting* method.

The crucial question is, of course, which branches are important enough to be retained for OTU picking. For sure, there is no point in retaining branches whose loss is negligible and comparable to that of most other branches. Such branches do not establish any recognizable pattern and hence they are not considered structurally important. I suggest two criteria to determine the breakpoint in an increasing series of losses. Figure 1 shows what may be a typical loss–rank plot. There are two points where the series distinctly changes its behavior: (1) the 7th point from the right follows immediately after the first big step in the series, i.e. after the first difference between successive values, which is considerably larger than usual; (2) the 11th point from the right follows immediately the place where the series forms an “elbow”, i.e. where the slow initial growth turns into fast final growth. It remains to determine which steps are larger than usual and where exactly the elbow lies. Currently, the “step” criterion is implemented as follows: (1) probability density distribution of step sizes (=differences between successive losses) is approximated by its kernel density estimate with bandwidth equal to the mean step size and rectangular kernel; (2) approximately zero probability density is defined as its 0.0001 quantile; (3) the intervals where probability densities drop at or below approximate zero are identified; and (4) the step sizes following the first such intervals are considered as outliers. Consequently, the first branch whose loss is separated from its predecessor in the ordered series by an outlying step is considered as the breakpoint (Figures 1, 2). The “elbow” criterion is defined by a simple geometric argument: in the loss–rank plot the first and the last loss is linked by a line and the point most distant from the line is the elbow and the next point is the breakpoint (Figure 1). The “elbow” criterion is therefore inherently more splitting compared to the “step” criterion.

**Figure 1.**
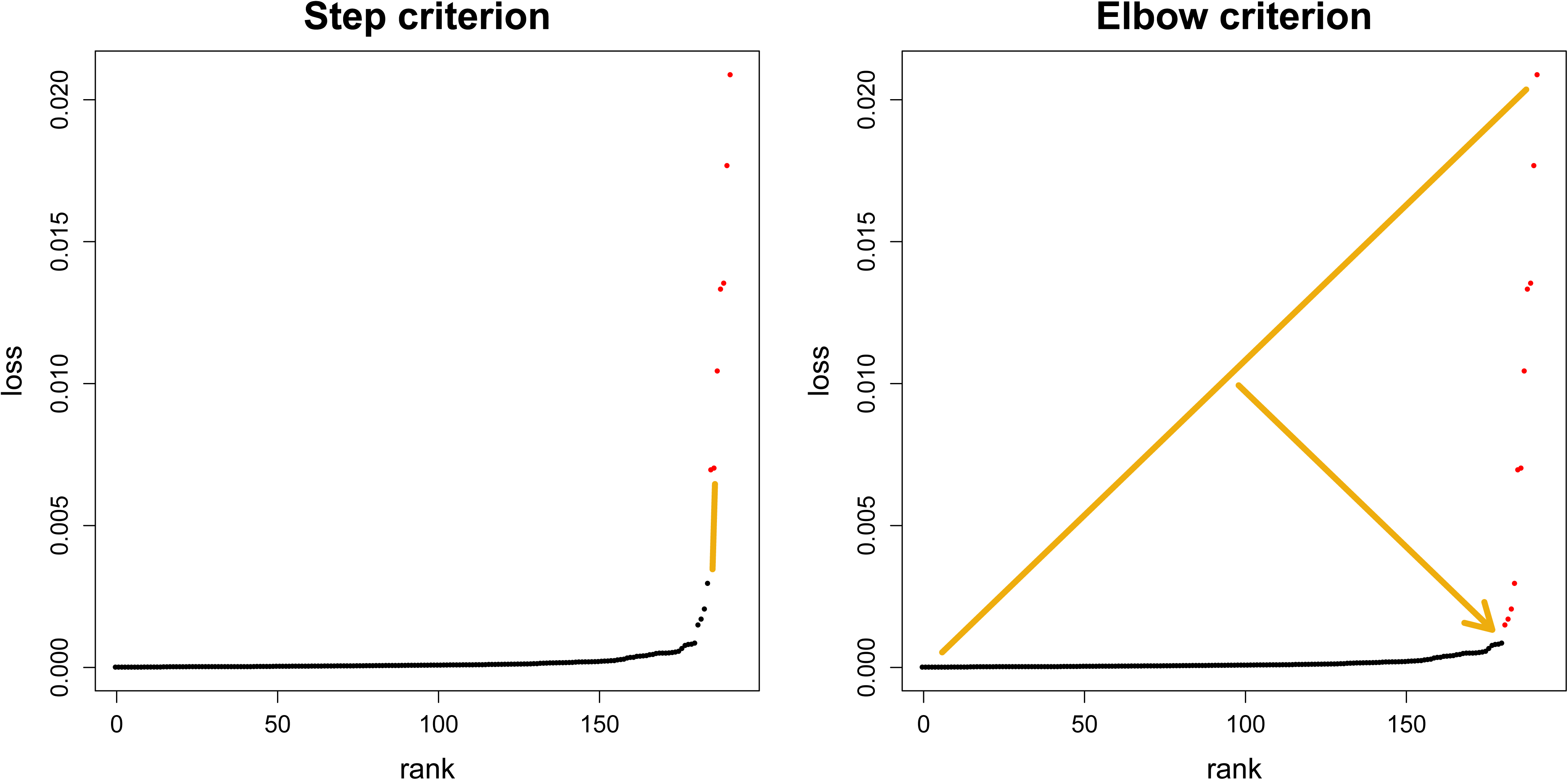
Step and elbow criterion as defined on the ordered loss series. Retained branches have ranks higher or equal to the first branch after any outlying difference between consecutive losses (step criterion) or after the first rank, where initial grow in loss suddenly accelerates (elbow criterion).

**Figure 2.**
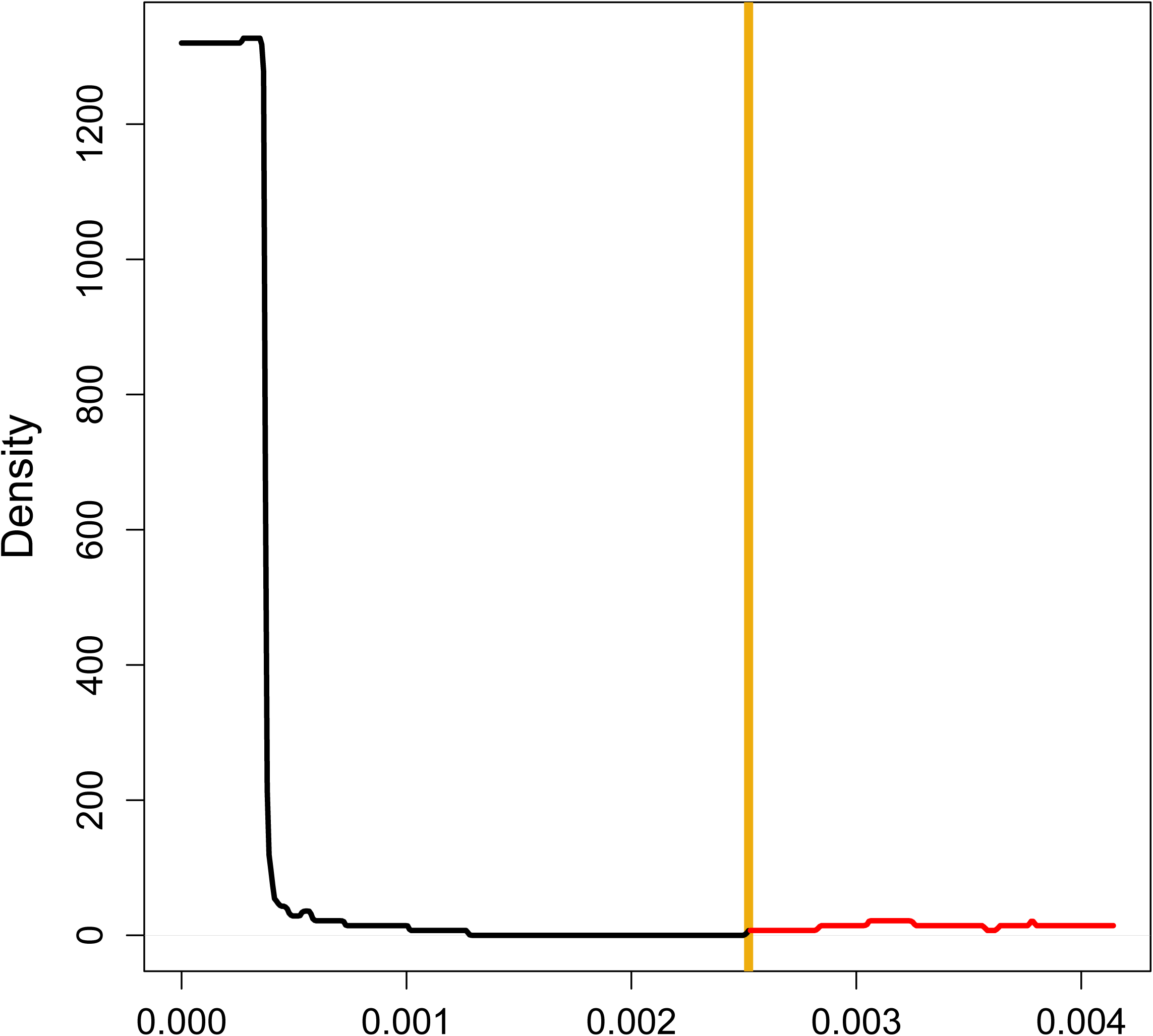
Kernel density estimate of step size distribution. Steps are differences between consecutive branch losses in their ordered series. The dark yellow line shows the threshold separating the first peak on the left (in black) from the rest (in red). The first step on the right from the threshold defines the step criterion.

### Multiple phylogenetic scales

Importantly, the method implicitly assumes the observed branching pattern is an overprint of just two processes, one consisting of much rarer events than the latter. This is likely true in many real cases, but not universally. As a consequence, the unique branches may be present at multiple, finer or coarser, scales. For instance, the tree may cover history of two radiations (each with several species) separated by a deep phylogenetic divergence, with every species represented by a sample of sequences whose relations are well described by the coalescent model. When applied to the tree as a whole, the branch-cutting method may recognize uniqueness of the branch separating the radiations, when applied to a subtree comprising just a single radiation it may recognize uniqueness of inter-specific branching and when applied to one intra-specific sample it may even recognize uniqueness of a deep coalescence event. Thus, the suggested criteria serve to recognize whether there is any recognizable pattern of unique branching present, but do not guarantee the pattern to be recognized at any particular resolution. The algorithm must be constrained to be directed to the resolution of interest. When it is informed which pairs of tree tips are to be considered heterospecific and/or conspecific, it can shift the estimated breakpoint to the closest value compatible with the constraints. In such case, the loss serves to leverage information available for a limited subset of data so it can be applied across the tree.

### Robustness

The branch-cutting algorithm retains branches whose removal significantly reduces average tip-to-tip distance. It follows it tends to retain branches that are long and central (basal in rooted trees) and discard short and peripheral (terminal) ones. Nevertheless, the central (basal) position promotes, but not guarantee, structural importance. There can be short central or basal branches that are not retained and discarding them may create OTU that is not monophyletic on the original tree. Delimitation of such OTU would be logically inconsistent and hence the relevant internal branches are retained too by the algorithm, in spite of non-conforming the formal (“step” or “elbow”) criterion.

Although long branches cannot make OTU delimitation logically inconsistent, they can more easily represent noise and/or sampling artifact rather than true genealogical separation. The branch-cutting method implicitly assumes phylogeny to be comprehensive and the intraspecific sampling to be equally intensive in every species. If some species are missing, long internal branches may dominate the branching pattern, even though they reflect sampling rather than branching process. If some species are poorly sampled, deep coalescence can be misidentified for interspecific differentiation and a couple of closely related alleles may erroneously appear as a true sample of distinct species. It may be also hard to detect rare and genetically uniform species that are likely represented by a single sequence, which bears no information about contrasting pace of speciation and coalescence process.

Finally, the algorithm assumes all branches are highly supported and their lengths are estimated with high confidence. Of course, this is not always true and poorly supported branches can mislead the algorithm, especially if they are long. Note, that the length of poorly supported branches is likely estimated with low confidence. This problem can be alleviated by *ad hoc* removal of branches whose support do not exceed given threshold or using trees containing polytomies in place of poorly supported branches.

### Implementation

The branch-cutting method was implemented in the form of package branchcutting for R (R Core Team 2018), which is freely available through CRAN (…) and offers the core ‘branchcutting’ function along with associated summary and plotting methods.

## Empirical examples

Usefulness of the branch-cutting method is demonstrated on four real mammal phylogenies inferred from published sequences of *cytochrome b* (*CYTB*) gene and mitochondrial control region (ctrl), which are documented in the Supplement 1. The organisms involved were Eurasian hedgehogs (*Erinaceus*), wood mouse (*Apodemus sylvaticus*), spiny mice (*Acomys*) and giraffes (*Giraffa*). All available haplotypes were re-analyzed using Bayesian inference in MrBayes 3.2.6 (Ronquist et al. 2012). HKY+G was used as the nucleotide substitution model. The alignment was partitioned according to locus (CYTB vs. ctrl, if the latter was sequenced) and in CYTB also according to codon position (12+3 scheme) and substitution model parameters (including overall substitution rates) were assumed to vary between partitions. Two analyses of each data set were conducted, one estimating unrooted tree with unconstrained branch lengths and the other estimating ultrametric tree based on strict clock assumption. Both analyses were run four times to check for convergence of Markov Chain Monte Carlo sampling from the posterior distribution. The posterior samples of both unrooted and ultrametric trees were represented by 50% majority consensus trees and the maximum clade credibility (MCC) trees. Every phylogeny was thus represented by four trees. The consensus trees were calculated in MrBayes and the clades with posterior probability < 0.10 were omitted which caused them to contain polytomies. The MCC trees were calculated using R packages ape (Paradis et al. 2004), phangorn (Schliep 2011) and phytools (Revell 2012). The reanalyzed data were of different size (59 individuals in *Erinaceus*, 63 in *Giraffa*, 317 in *Acomys* and 517 in *A. sylvaticus*) and they differed also in their phylogenetic extent: while only single taxonomic species is assumed to be included in *A. sylvaticus*, there may be about four of them in *Erinaceus* and *Giraffa* and up to about twenty in *Acomys*.

### Results of the branch-cutting

Four species of *Erinaceus* hedgehogs are currently recognized (Hutterer 2005) and their distribution serves as a classical example of biogeographic pattern induced by the last continental glaciation (Hewitt 1999). Not only that *E. concolor, E. roumanicus* and *E. europaeus* are separated by dispersal barriers or forming stable contact zones, but also the intraspecific phylogenetic structure of *E. europaeus* is suggestive of the past fragmentation in different Mediterranean refugia (Seddon et al. 2001, 2002; Bolfíková and Hulva 2012). The fourth species, *E. amurensis*, lives quite far apart, in the basins of Amur and Yangtze Rivers (Cassola 2016). In three out of four trees the step criterion identified six OTUs exactly corresponding to the recognized species and phylogeographic lineages of *E. europaeus* (Figure 3). Only in the unrooted consensus tree two OTUs were identified, one corresponding to *E. concolor* + *E*.*roumanicus* and the other to the rest. Even in this case, however, the distribution of differences in branch-specific losses was close to support six OTUs instead of two, which is apparent from its similarity to the distribution based on the unrooted MCC tree (Figure 4). Building upon the prevailing six-OTU solution I examined the effect of assuming all *E. europaeus* individuals to be conspecific. Using this constraint the four recognized species were recovered from the ultrametric trees, while in the unrooted MCC tree also *E. concolor* and *E.roumanicus* were merged (Figure S2.1, i.e. Fig. 1 in the Supplement 2). In fact, divergence between the latter species is of comparable scale to *E. europaeus* lineages, but at least in the ultrametric trees branches supporting them appeared less redundant in the overall picture of the tree than the branches constrained to be encompassed by *E. europaeus*. The elbow criterion recovered seven to eleven OTUs in the hedgehog trees (Figure S2.2).

**Figure 3.**
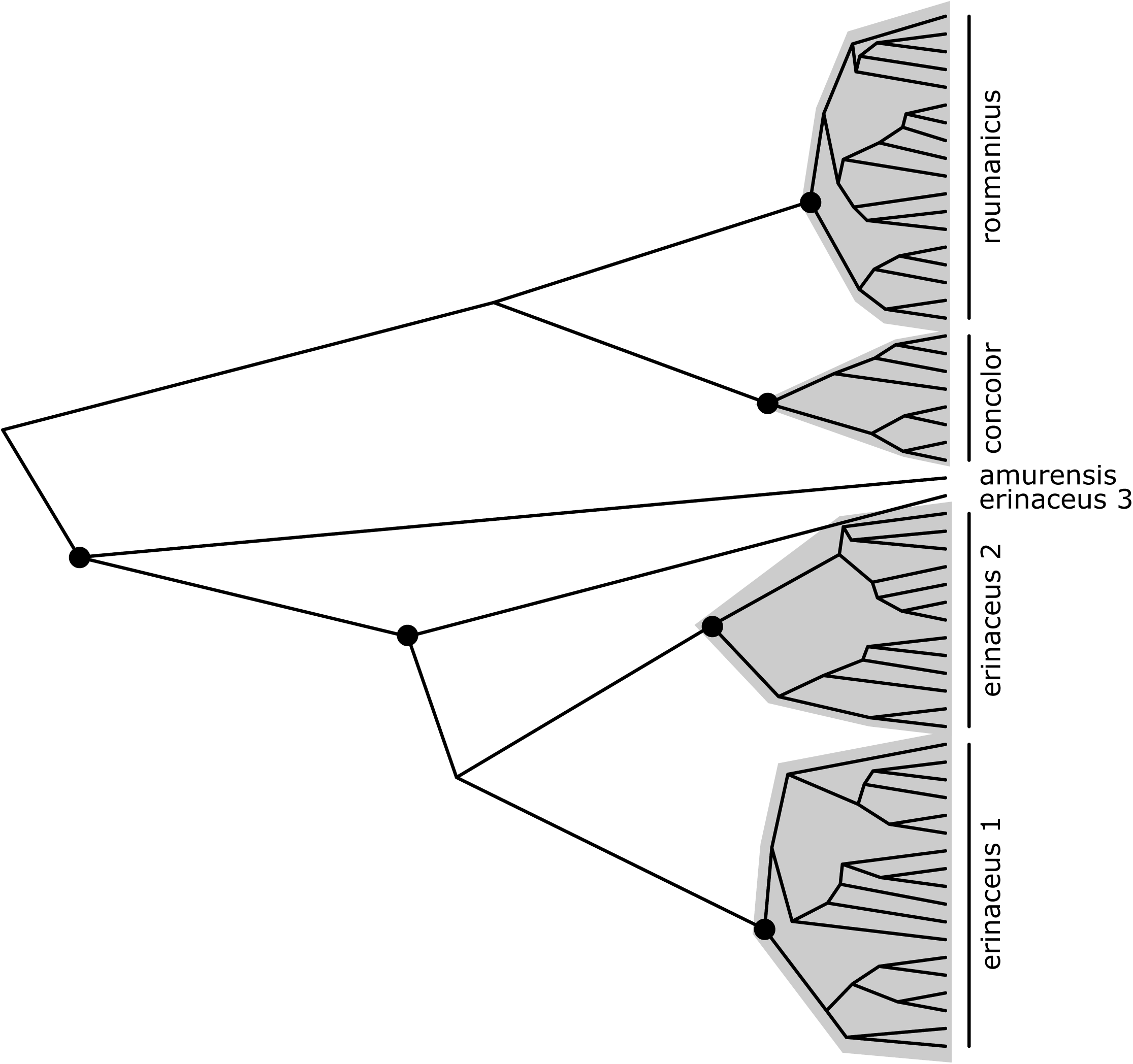
Six OTUs delimited on the ultrametric MCC tree of *Erinaceus* according to the unconstrained step criterion. Parts of the tree corresponding to particular OTUs are shaded in grey, dots at the tree nodes mark the most recent common ancestors of the OTUs.

**Figure 4.**
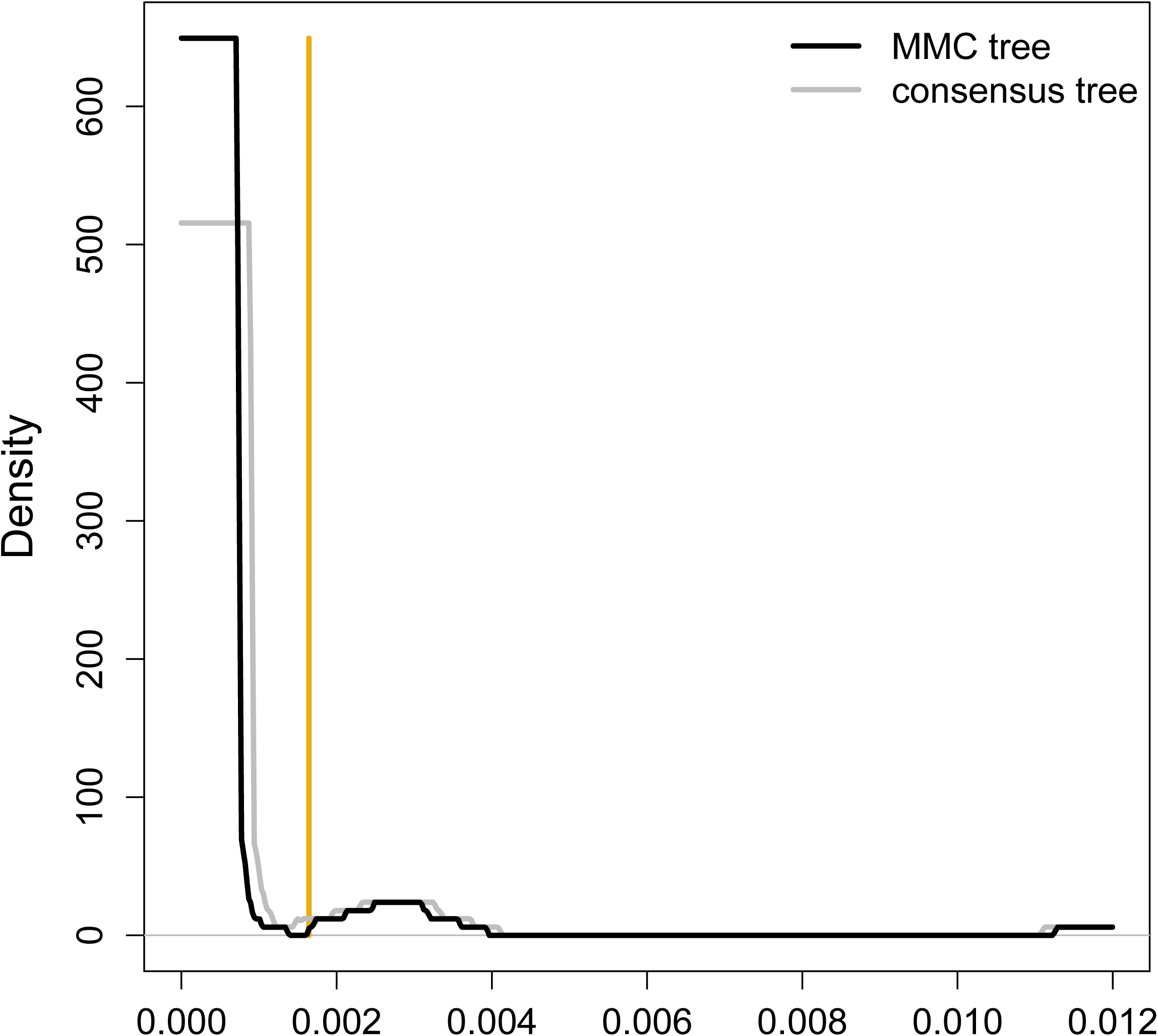
Comparison of step size density distributions between unrooted MCC and consensus trees of hedgehogs, showing they are essentially identical and delimitation of two instead of six OTUs on the latter is due to minute differences in branch length distribution in the tree. The dark yellow line shows the threshold based on the unrooted MCC tree.

The phylogeographic structure of *A. sylvaticus* also bears imprint of glacial cycles. In their comprehensive study of CYTB variation Herman et al. (2017) identified six phylogeographic lineages, three corresponding to the formerly recognized Mediterranean refugia (Michaux et al. 2003), one comprising North-African populations, one being detected on the Channel Islands and the last one (‘peripheral’) being tentatively interpreted as the remnant of the first wave of post-glacial recolonization. All these populations are unanimously considered conspecific (Musser & Carleton 2005, Wilson et al. 2017). The step criterion identified eight to twenty OTUs here. Both consensus trees supported eight OTUs that were much better interpretable compared to finer delimitation recovered from the MCC trees (Figure S2.3). Thus, having polytomies in place of poorly supported branches proved beneficial in this case. Interestingly, however, the OTUs from the two consensus trees were not identical. The ultrametric consensus supported pooling of the geographically adjacent ‘Sicilian’ and ‘south-eastern’ lineages and partition of the ‘central’ and ‘peripheral’ lineages into geographically meaningful, yet overlapping sublineages. The unrooted consensus supported solution that was nearly identical to Herman et al.’s lineages, only two haplotypes stood slightly aside their most closely related OTUs: ‘south-eastern’ and ‘peripheral’, respectively (Figure 5). Given their phylogenetic proximity, it was no surprise that constraining them to be conspecific with their nearest neighbors had no effect elsewhere in the tree and the six-OTU delimitation of Herman et al. was recovered. The elbow criterion suggested very many (18–159) OTUs here.

**Figure 5.**
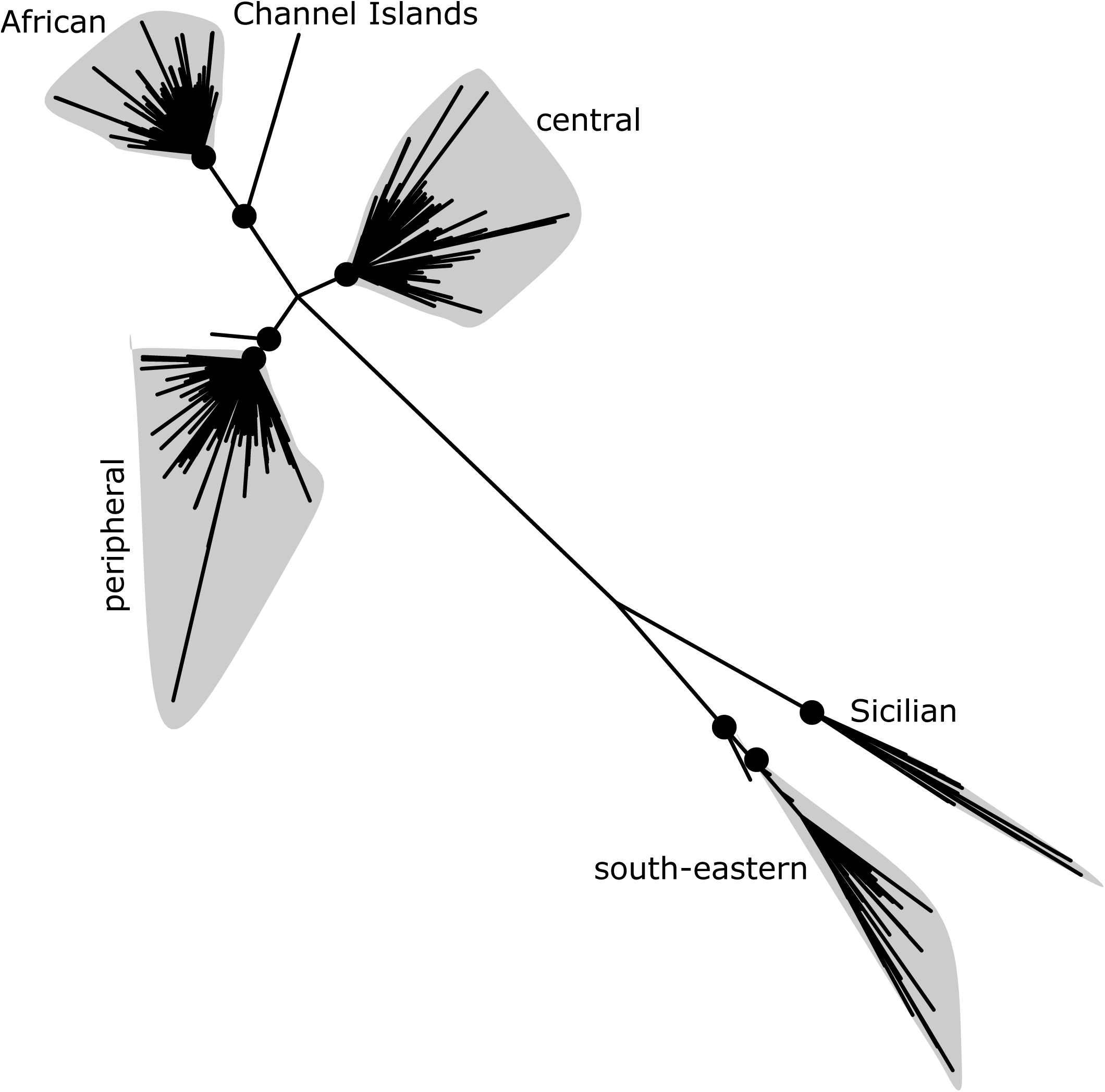
Phylogenetic proximity of OTUs delimited on the unrooted consensus tree of *A. sylvaticus*. Two of the OTUs were very small and both were found very close to some other OTUs as compared to other inter-OTU distances. That is where the conspecific constraints were applied.

The recently published phylogeny of *Acomys* (Aghová et al. in review) is the first one covering the whole genus, which is distributed in the sub-Saharan Africa, Middle East and part of the Mediterranean. The most recent compendia recognize about 20 species (Musser and Carleton 2005, Wilson et al. 2017), while the provisional classification by Aghová et al. recognizes up to 26 OTUs in seven major phylogenetic lineages. The branch-cutting with the step criterion recovered almost identical delimitations in all four trees, with fourteen or fifteen OTUs in total. There were only two small groups of individuals that were variously split in some trees but not in others and constraining both of them to keep together resulted in twelve OTUs (Figure 6), which is arguably the most robust delimitation currently available for the genus. The elbow criterion delimited from 38 to 42 OTUs.

**Figure 6.**
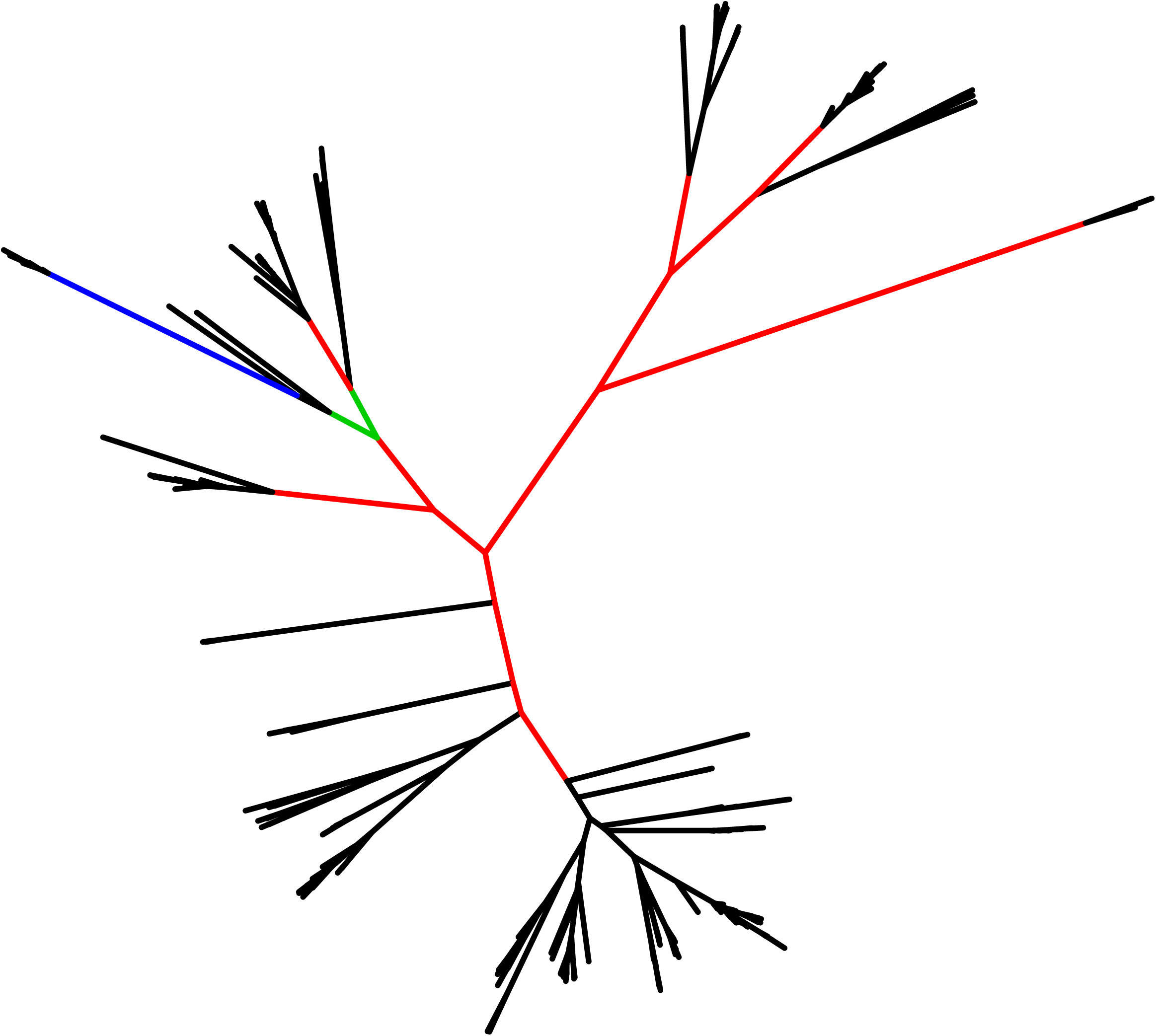
Unrooted MCC tree of *Acomys* with twelve OTUs conforming the conspecific constraints. The colors indicate branches to be collapsed (black), retained (red), retained due to position in the tree in spite of not passing the step criterion (green) and those that are to be collapsed just to preserve monophyly of OTUs, whose delimitation was enforced by conspecific constraints (blue).

Although the giraffe was traditionally recognized as monotypic with multiple subspecies (Grubb 2005, Wilson et al. 2011, Bock et al. 2014), recently it was suggested some subspecies are not only differentiated in their fur coloration, but also genetically distinct and hence deserve species status. The re-analysis is based on data presented by Fennessy et al. (2016) who analyzed *CYTB* and ctrl mitochondrial loci as well as seven nuclear markers and concluded there are four giraffe species robustly supported by nuclear data, although the mitochondrial tree alone would suggest presence of seven lineages, designated subspecies by the authors. The step criterion basically suggested four OTUs to be present. In one case (unrooted consensus tree) five OTUs were recovered but two of them were phylogenetically very close and they were considered distinct just to preserve monophyletic condition (Figure S2.4). In other trees (and also the nuclear tree) they were monophyletic and their separation in the unrooted consensus likely stems from the tree imperfection. Interestingly, however, four OTUs delimited here were not concordant with four nuclear species of Fennessy et al. (2016). The nuclear species *G.giraffa* was split in two OTUs corresponding to mitochondrial subspecies *giraffa* and *angolensis*, whose divergence was already noted (Bock et al. 2014), but here they were even not monophyletic (Figure 7). On the contrary, the nuclear species *G. camelopardalis* and *G.reticulata* were pooled into a single OTU. When *G.reticulata* was constrained to be heterospecific from *G.camelopardalis*, the branch-cutting method identified seven or eight OTUs (on unrooted and ultrametric trees, respectively, Figure S2.5). These OTUs corresponded to the subspecies of Fennessy et al. (2016), with the subspecies *tippelskirchi* possibly split in two. Analyses based on the elbow criterion returned from five to nine OTUs.

**Figure 7.**
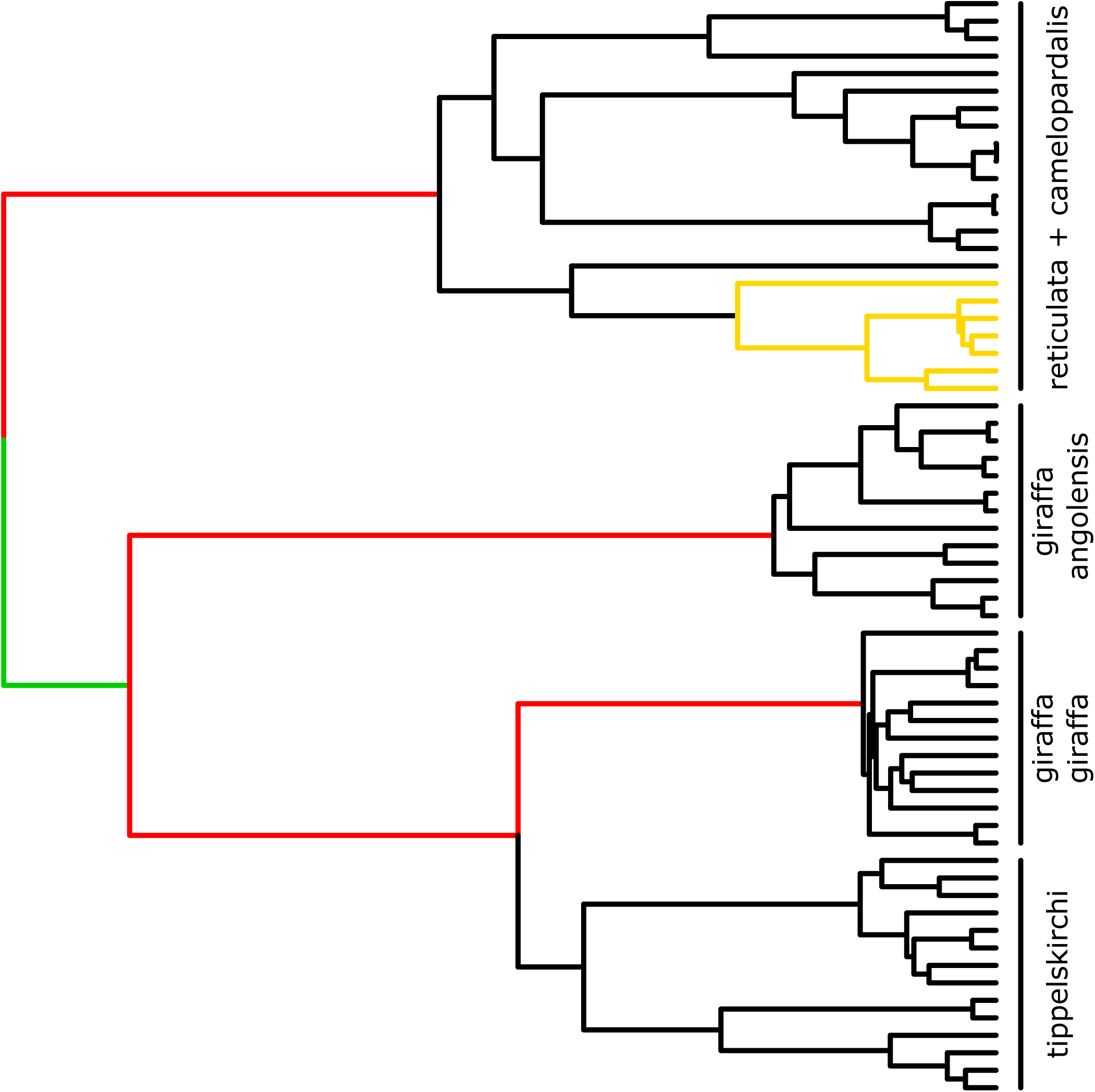
Ultrametric MCC tree of *Giraffa* with four OTUs and the position of *G. reticulata* delimited by nuclear genes indicated. *G. reticulata* is shown in gold, otherwise the same color code is used as in Figure 6.

### Comparison to other methods

Table 1 summarizes number of OTUs identified by branch-cutting and three other methods operating on single locus data. *The Generalized Mixed Yule Coalescent* (GMYC) method operates on a fully bifurcating, rooted and ultrametric tree. In its basic version (Pons et al. 2006, Fujisawa et al. 2013) it uses maximum likelihood to estimate a single threshold time where all delimited species originate and which classifies branching times in two distributions, whose shapes conform expectations of the constant rate pure speciation process (Yule 1924; Nee 1994) and the neutral coalescent (Kingman 1982; Hudson 1990). In its multi-threshold version (mGMYC) it allows delimited species to differ in their time of origin (Monaghan et al. 2009). The *Poisson Tree Processes* (PTP) method uses fully bifurcating, rooted trees, which may not be ultrametric, however. It assumes branch lengths to differ according to whether they are affected by speciation dynamics or just mutation and drift and the two classes of branches are bounded by switch points on a tree, whose locations are inferred using maximum likelihood (Zhang et al. 2013). Recently, its multi-rate extension (mPTP) was devised, allowing for lineage-specific distributions of intraspecific branch lengths (Kapli et al. 2017). Finally, *Automatic Barcode Gap Discovery* (ABGD; Puillandre et al. 2012) aims to identify a unique ‘barcode gap’: genetic distance threshold allowing to decide, whether any two individuals are conspecific or not. It therefore operates on a matrix of pairwise distances.

**Table 1.**
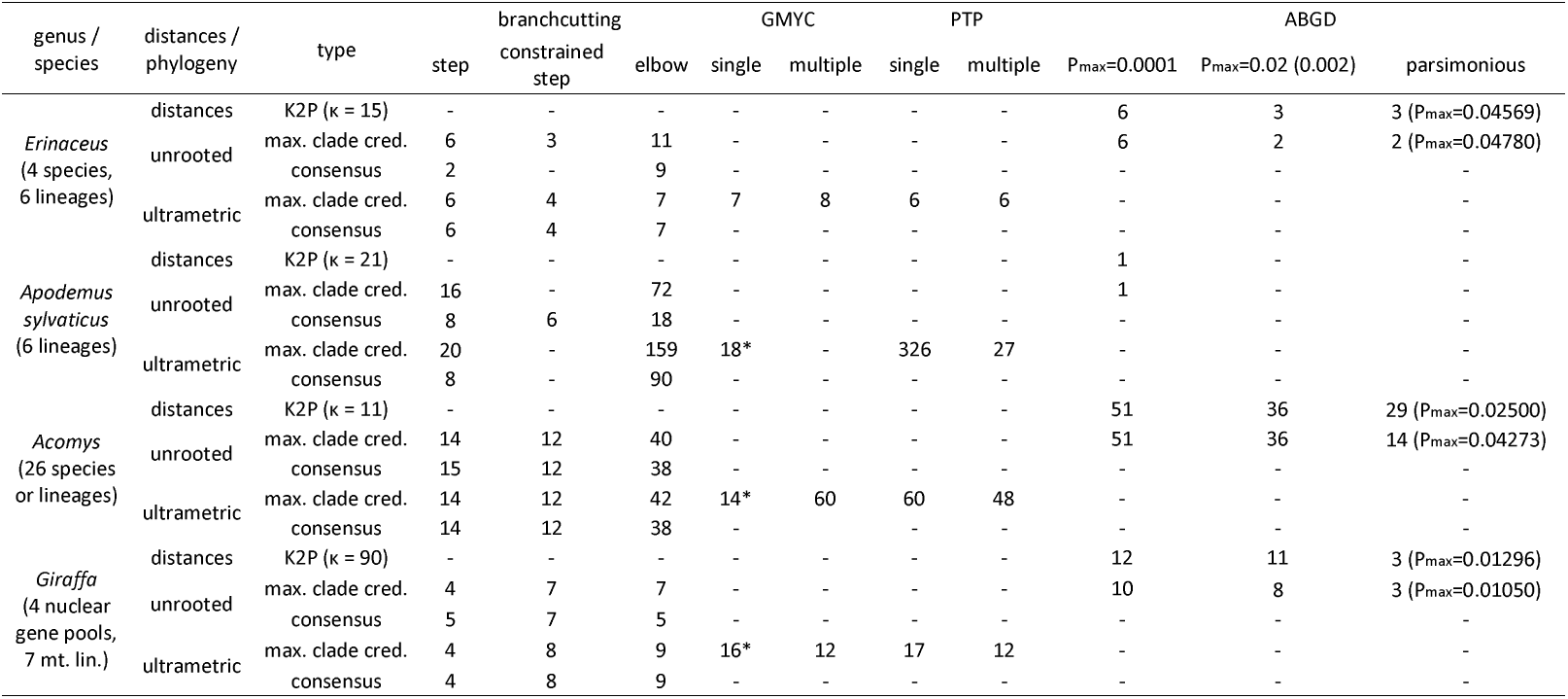
Number of OTUs delimited by different OTU-picking methods. The cells are empty if the method is not applicable to the particular type of data or if there was no point in application of the branchcutting constraints. Asterisks denote GMYC solutions based on the first likelihood peak rather than on the global likelihood maximum.

GMYC/mGMYC and PTP/mPTP analyses were performed on ultrametric MCC trees, although PTP/mPTP would be applicable also for non-ultrametric trees. The consensus trees could not be used as they contained polytomies. ABGD analysis used matrix of either Kimura two-parameter distances (Kimura 1980) or phylogenetic distances observed on unrooted MCC trees. Optimization of scaling parameters in the GMYC/mGMYC models was integral to model fitting, while two tuning parameters of ABGD (maximum intraspecific distance, *P*_max_, and relative gap width, *X*) had to be specified prior to analysis. Three different values of *P*_max_ were used in turn: 0.0001 motivated by the default collapse height parameter in STACEY (Jones et al. 2015), 0.02 motivated by results of Bradley and Baker (2001) and the largest *P* suggesting more than one OTU (“most parsimonious delimitation”). *X* was set to 1.5. GMYC and mGMYC models were fit in R package ‘splits’ (Ezard et al. 2017), PTP and mPTP models via web-service (http://mptp.h-its.org) and ABGD solution was also found using on-line application (http://wwwabi.snv.jussieu.fr/public/abgd/abgdweb.html).

The numbers of delimited OTUs varied widely, but usually they were larger than those obtained by the branch-cutting method. The hedgehog data set stood out of this observation as the six-OTU delimitation (recognized species + phylogeographic lineages) was recovered also by PTP, mPTP and ABGD with *P*_max_=0.0001. In the other three data sets (besides the hedgehogs) GMYC likelihood profiles showed more or less clearly pronounced peak at some threshold time, followed by local minimum and then by steady increase continuing until the threshold time of zero or nearly so (Figure S2.6). On the other hand, delimitations corresponding to the first peak were roughly comparable to those of other methods. mGMYC might output either lower or higher number of OTUs and in the case of *A.sylvaticus* it ended with error, but there is actually a warning given by the authors of the package stating the implementation is experimental. PTP tended to output apparently excessive numbers of OTUs, but this problem was largely alleviated by application of mPTP. Finally, ABGD solutions depended much on *P*_max_ parameter. If it was low (0.0001), directing the algorithm to fine phylogenetic scale, it might return results congruent with other biological evidence (like in *Erinaceus* or *A.sylvaticus*), but also to hugely oversplit the taxa (like in *Giraffa* and especially *Acomys*). Shift at an intermediate level (set to be 0.02 in small mammals, but 0.002 in *Giraffa*) might or might not have substantial effect and perhaps the safest strategy to get some biologically interpretable solution was to increase *P*_max_ as long as it allowed more than one OTU to be delimited.

## Discussion

It is now widely acknowledged speciation is not an instantaneous event but a protracted and complex process (Rosenblum et al. 2012; Dynesius and Jansson 2014; Etienne et al. 2014). Even more, the very concept of species is time scale dependent: populations that behave as perfectly distinct species on a shorter time scale (say 1000 generations) may eventually merge and won’t be recognized as distinct on a longer time scale (say 100,000 or 1 million generations). A single gene tree generally provides limited insight into this complexity and thus it is not expected to reliably resolve species limits (Yang and Rannala 2010). Its analysis has a great heuristic value, however. First, clustering of its tips provides information about differentiation of that particular gene, which is often biologically interpretable. Typical example here is mitochondrial phylogeography, where the lower effective population size and possibly also lower female dispersal causes distinctive clusters of sequences to be indicative of spatially coherent populations. Second, provisionally delimited units can be used in downstream analyses, that are likely robust to a degree to imprecise classification. This is the case for analyses of biodiversity and its geographic patterns. Finally, OTUs can be considered as candidate species, i.e. working hypotheses about species boundaries that are to be refined by other data and analyses (e.g. by multispecies coalescent analysis of multilocus nuclear data).

The branch-cutting method introduced here is a novel OTU picking method from the ‘pattern-recognition’ family, which means it makes no parametric assumptions about the evolutionary process. Instead, it merely assumes the observed phylogenetic differentiation is an overprint of different processes and targets signature of their distinct dynamics. The above examples show the assumption is reasonable enough to provide basis for delimitation of biologically meaningful units. In the hedgehog data set it recognizes exactly the same units as delimited previously on the basis of existing biological knowledge (Seddon et al. 2001, 2002) and within *A. sylvaticus* it also approaches the phylogeographical delimitation (Herman et al. 2017). In *Acomys* example it appears to sort out “nuts and bolts” as it returns delimitation supporting the best evidenced species boundaries (e.g. *A. muzei* × *A.ngurui*; Verheyen et al. 2011; Petružela et al. 2018), lumping those whose separate status was deemed doubtful (e.g. Mediterranean endemics; Giagia-Athanasopoulou et al. 2011) and suggesting novel taxa in yet other parts of the tree (namely in the *wilsoni* group of species). Finally, analysis of the giraffe data set shows how useful branch-cutting can be in formulation of explicit hypotheses. There is discrepancy in delimitation based on nuclear and mitochondrial genes and implicit in the taxonomy of Fennessy et al. (2016) is the hypothesis the mitochondrial lineages represent finer-scale resolution perhaps with some genealogical discrepancy due to incomplete lineage sorting. The branch-cutting delimitation shows, however, that some inter-subspecific mitochondrial divergences are considerably deeper than inter-specific ones. Although this may be still explained by incomplete lineage sorting and/or introgression (see discussion in Bock et al. 2014), there is also an alternative hypothesis that the nuclear species *G. reticulata* was supported as such only due to limited sampling. If it was the case, there would be only three nuclear species left: *G. giraffa* encompassing deep mitochondrial divergence, *G. camelopardalis* encompassing shallow mitochondrial divergence and *G.tippelskirchi* corresponding to the remaining mitochondrial OTU.

The examples above also suggest some caution when interpreting branch-cutting delimitations as they prove to be somewhat different when based on different trees inferred from the same sample of sequences. Even if topology is identical across the trees, relative branch lengths depend on the model (e.g. clock vs. non-clock) and the summary method used and they are always estimated with some error. In some cases a moderate difference in the length of particular branch may become a tipping point deciding which branches are retained for the OTU delimitation. This difference is expected to be especially pronounced between fully bifurcating trees and those containing polytomies and while omission of poorly supported branches may be beneficial in some cases (c.f. *A. sylvaticus*), it may be detrimental in others, where too severe collapsing of branches leads to large loss of information. The difference in distribution of branch lengths and hence in loss score may also blur distinction between large and small steps in the ordered loss series and the step criterion may miss the step that is otherwise detected as outlying. This apparently happened in the case of *Erinaceus* unrooted consensus tree, where only one branch is retained, in contrast to the unrooted MCC tree with very similar loss-rank profile, but seven retained branches.

The elbow criterion always retains at least the same number of OTUs as the step criterion, but usually (much) more. It appears to be self-similarity of branch length distribution what causes the elbow to correspond to still lower ranked losses as the size of phylogeny grows. It is then no surprise that phylogeny of *A. sylvaticus* with 517 tips has up to 159 OTUs according to the elbow criterion. Thus, it is the step criterion which is pivotal for the branch-cutting method. Nonetheless, the elbow criterion may be still useful under some circumstances. A possible application is delimitation of candidate species to be investigated in a subsequent multispecies coalescent analysis, e.g. in BPP (Yang and Rannala 2010) or STACEY (Jones et al. 2015). Given the elbow criterion is likely to oversplit the taxon, its OTUs may represent a safe starting point for subsequent merging, while providing considerable simplification of the problem.

In comparison with other single-locus methods, two points are worth to be stressed. First, branch-cutting appears to be much less prone to oversplitting than any other alternative. Table 1 shows it usually suggested lower (and sometimes much lower) numbers of OTUs than the alternatives and even more importantly its OTUs were often well congruent with expectations based on external information. Second, branch-cutting is very versatile, allowing to work with any phylogenetic tree, rooted or unrooted, bifurcating or not and having branch lengths in any units. It could even operate on reticulated networks instead of trees, although I did not explore this possibility here. It also makes generally weaker assumptions about the nature of the underlying evolutionary process, especially compared to the model-based methods. On the other hand, of course, GMYC and PTP models allow the corresponding methods to operate in the likelihood framework and enjoy its power including rigorous model selection and assessment of confidence limits. In addition, it is also good to note that branch-cutting may be more demanding with respect to sampling. Unbalanced sampling may left long internal branches within a genealogically homogenous species and this one can be therefore split into several OTUs. This is less likely in PTP/mPTP as they model branch lengths by a mixture of exponential distributions whose long tails could easily account for an excessive branch. In contrast, singletons (single representatives of their species) may not be recognized as such by the branch-cutting. The more individuals are sampled from a species the more pairwise distances are affected by removal of its stem branch and the single terminal branch establishing a singleton have to be quite long to be retained. This is much lesser problem for GMYC/mGMYC due to their reliance on branching times instead of branch lengths.

Anyway, all the methods compared here revolve around the key assumption of a diagnostic difference between intraspecific and interspecific variation. In reality, however, more than two processes can shape the tree, e.g. constant rate diversification might be punctuated by a rapid radiation within one of the clades and recurrent population splits and mergers might take place within the species. In this respect Darwin’s finches may serve as an illustrative example (Grant and Grant 2007). Such complexity can blur distinction between traces of speciation/extinction and mutation/drift dynamics and eventually make them inseparable. OTU-picking methods differ in the ways they approach the problem. GMYC-based and PTP-based methods, as currently implemented, assume speciation process to leave a single imprint on the tree and rely on their parametric assumptions and maximum likelihood fitting procedure to distinguish it from intra-specific patterns. Their assumptions may include, however, species-specific threshold times (mGMYC) or branching rates (mPTP). AGBD acknowledges multiple barcoding gaps may be present and requires two parameters to direct the algorithm to the gap, deemed to separate inter-specific and intra-specific variation. Branch-cutting requires no parameters to be specified a priori, but it identifies any branch of outlying structural importance, no matter what evolutionary process it actually represents. It does not involve any tuning aiming to identify phylogenetic scale of ‘true’ speciation events, but instead it allows to use external information about the nature of differentiation patterns by means of conspecific and/or heterospecific constraints. Given the time scale dependency of species as entities, there is no firm distinction between temporary structures created, for instance, by ongoing geographical isolation and permanent structures that arose due to irreversible reproductive isolation. This resignation on determination of the ‘true’ species from the sequence variation alone echoes what was proclaimed about multispecies coalescent methods: they delimit “structure, not species” (Sukumaran and Knowles 2017).

In any case it is advisable to critically consider the branch-cutting solutions and if there are any issues, address them by repeating the analysis with meaningful constraints. The constraints are applied to the ordered loss series and they may therefore affect also other comparably differentiated OTUs. Two conspecific constraints applied to *Acomys* may serve as a good example. Crucial here was the mutual comparison of OTUs from analyses based on different trees which delimited up to 15 OTUs. Closer inspection revealed there are two pairs of OTUs that are either merged or split, depending on the particular tree used. Imposing conspecific constraints on both of them should reduce the number of OTUs to 13, but in fact twelve OTUs were returned because the constraints caused yet another OTU to merge with one of the pairs. In contrast, constraining outlying haplotypes of *A. sylvaticus* to be conspecific with their nearest neighbors in the phylogeny caused no other merger and the delimitation proposed in the original publication was returned. These constraints were not motivated only by congruency with the published delimitation, however. The very pattern of a few haplotypes standing separately but much closer to their neighboring OTUs than otherwise observed (Figure 5) was suggestive of delimitation that inappropriately considers deep coalescent branch as structurally important.

Development of the branch-cutting method was motivated by a necessity to consistently delimit taxonomic units in phylogeographic and systematic studies, either as entities of interest or as candidate species whose distinctiveness is to be tested by further collection of data and application of more sophisticated analyses. At least in this context branch-cutting appears to be a powerful alternative to the existing OTU-picking methods. It could be employed, however, also in barcoding and meta-barcoding studies with their large phylogenies. It would probably require thoughtful definition of heterospecific constraints and cutting of interspecific branches prior to the analysis, so the algorithm would focus on differentiation patterns typical for shallow parts of the tree.

## Acknowledgements

Financial support for this study was provided by the Czech Science Foundation, project no. 18-17398S. I thank to J. Bryja for commenting the first version of the manuscript.

